# Reciprocal carbon subsidies between autotrophs and bacteria in stream food webs under stoichiometric constraints

**DOI:** 10.1101/447987

**Authors:** Benoît O.L. Demars, Nikolai Friberg, Joanna L. Kemp, Barry Thornton

**Affiliations:** Norwegian Institute for Water Research, Gaustaallen 21, 0349 Oslo, Norway; The James Hutton Institute, Craigiebuckler, Aberdeen AB15 8QH, UK; University of Copenhagen, Freshwater Biological Section, Universitetsparken 4, 3rd floor, 2100 Copenhagen, Denmark; University of Leeds, water@leeds, School of Geography, Leeds LS2 9JT UK

**Keywords:** reciprocal subsidies, microbial loop, dissolved organic matter, nutrient spiralling, whole-stream metabolism, flow food web

## Abstract

1. Soils are currently leaching out their organic matter at an increasing pace and darkening aquatic ecosystems due to climate and land use change, or recovery from acidification. The implications for stream biogeochemistry and food webs remain largely unknown, notably the metabolic balance (biotic CO_2_ emissions), reciprocal subsidies between autotrophs and bacteria, and trophic transfer efficiencies.
2. We use a flow food web approach to test how a small addition of labile dissolved organic matter affects the strength and dynamics of the autotrophs-bacteria interaction in streams. Our paired streams whole-ecosystem experimental approach combined with continuous whole-stream metabolism and stable isotope probing allowed to unravel carbon fluxes in the control and treatment streams.
3. We increased the natural supply of dissolved organic matter for three weeks by only 12% by continuously adding 0.5 mg L^−1^ of sucrose with a δ^13^C signature different from the natural organic matter. Both photosynthesis and heterotrophic respiration increased rapidly following C addition, but this was short lived due to N and P stoichiometric constraints. The resulting peak in heterotrophic respiration was of similar magnitude to natural peaks in the control observed when soils were hydrologically connected to the streams and received soil derived carbon.
4. Carbon reciprocal subsidies between autotrophs and bacteria in the control stream accounted for about 50% of net primary production and 75% of bacterial production, under low flow conditions when stream water was hydrologically disconnected from soil water. The reciprocal subsidies were weaker by 33% (autotrophs to bacteria) and 55% (bacteria to autotrophs) in the treatment relative to the control. Net primary production relied partly (11% in the control) on natural allochthonous dissolved organic carbon via the CO_2_ produced by bacterial respiration.
5. Many large changes in ecosystem processes were observed in response to the sucrose addition. The light use efficiency of the autotrophs increased by 37%. Ecosystem respiration intensified by 70%, and the metabolic balance became relatively more negative, i.e. biotic CO_2_ emissions increased by 125%. Heterotrophic respiration and production increased by 89%, and this was reflected by a shorter (−40%) uptake length (Sw_OC_) and faster (+92%) mineralisation velocity of organic carbon. The proportion of DOC flux respired and organic carbon use efficiency by bacteria increased by 112%.
6. Macroinvertebrate consumer density increased by 72% due to sucrose addition and consumer production was 1.8 times higher in the treatment than in the control at the end of the experiment. The trophic transfer efficiencies from resources to consumers were similar between the control and the treatment (2-5%).
7. *Synthesis*. Part of the carbon derived from natural allochthonous organic matter can feed the autotrophs via the CO_2_ produced by stream bacterial respiration, intermingling the green and brown webs. The interaction between autotrophs and bacteria shifted from mutualism to competition with carbon addition under nutrient limitation (N, P) increasing biotic CO_2_ emissions. Without nutrient limitation, mutualism could be reinforced by a positive feedback loop, maintaining the same biotic CO_2_ emissions. A small increase in dissolved organic carbon supply from climate and land use change could have large effects on stream food web and biogeochemistry with implications for the global C cycle under stoichiometric constraints.

## Introduction

The global annual riverine flux of organic C (0.26-0.53 Pg C year^−1^) to the oceans is comparable to the annual C sequestration in soil (0.4 Pg C year^−1^), suggesting that terrestrially derived aquatic losses of organic C may contribute to regulating changes in soil organic carbon storage (Dawson 2013). Soils are currently leaching out their organic matter at faster rates to aquatic ecosystems due to climate and land use change, or recovery from acidification (e.g. Freeman et al. 2004, Monteith et al. 2007, Drake et al. 2015). The fate of this organic carbon in riverine systems remains poorly understood at the global scale, notably the degassing of CO_2_ back to the atmosphere (Drake et al. 2018), and remains highly debated regarding its contribution to aquatic food webs (e.g. Cole 2013, Brett et al. 2017).

The bacterial processing rate of terrestrial organic matter is influenced by the strength and complexity (aromaticity) of the carbon bonds of the organic matter and C:N:P ecological stoichiometry (Mineau et al. 2016, Evans et al. 2017, Kominoski et al. 2018). Numerous labile dissolved organic carbon (DOC) additions (from trace amount to 20 mg C L^−1^) have repeatedly shown DOC use by bacteria and its transfer through the food chain (e.g. Warren et al. 1964, Hall 1995, Hall and Meyer 1998, Parkyn et al. 2005, Wilcox et al. 2005, Augspurger et al. 2008). Fewer studies used leaf leachate material including less labile DOC (e.g. Cummins et al. 1972, Friberg and Winterbourn 1996, Wiegner et al. 2005, Wiegner et al. 2015). Many studies have also traced the flux of autochthonous DOC through to the bacteria (e.g. Lyon and Ziegler 2009, Risse-Buhl et al. 2012, Hotchkiss et al. 2014, Kuehn et al. 2014).

The interactions between autotrophs and bacteria are however difficult to study (Amin et al. 2015), because primary producers, decomposers and organic matter (allochthonous and autochthonous) are intricately connected both in benthic biofilms (Kamjunke et al. 2015, Battin et al. 2016) and pelagic aggregates (Grossart 2010). In theory, bacteria and autotrophs could compete for limiting nutrients (Currie and Kalff 1984), notably when bacteria have lower C:nutrient biomass ratios than autotrophs (Daufresne and Loreau 2001). However, bacteria can have high C:nutrient ratios similar to autotrophs (Cotner et al. 2010) supporting the co-existence of autotrophs and heterotrophs (Daufresne and Loreau 2001). Correlative analyses in streams seem to support the idea that positive interactions between autotrophs and bacteria increase with nutrient (N, P) limitation (Carr et al. 2005, Scott et al. 2008). Whole-stream metabolism in open streams under low flow conditions (hydrologically disconnected from catchment soils supplying allochthonous organic carbon) showed ecosystem respiration to be tightly correlated to gross primary production (Demars et al. 2016), suggesting a strong indirect mutualistic interaction between autotrophs and bacteria (Demars et al. 2011a), with autotrophs providing detritus C, N, P to bacteria and bacteria regenerating N and P by mineralisation (Cotner et al. 2010, Demars et al. 2011b), in agreement with theory (Daufresne and Loreau 2001).

The potential benefit of bacterial CO_2_ for primary producers has been hypothesised to explain a small increase (16-20%) in gross primary production following dissolved organic carbon addition and associated increase in bacterial activities (Robbins et al. 2017). The reciprocal carbon subsidies between autotrophs and bacteria in stream food webs has however, to our knowledge, not been studied either empirically or theoretically. It raises the possibility that the carbon of the primary producers may be partly derived from allochthonous organic matter processed by the bacteria, and intermingle the green and brown webs (Zou et al. 2016), an overlooked issue in the autochthony-allochthony debate (e.g. Cole 2013, Brett et al. 2017).

Here we test how dissolved organic carbon addition affects the strength and dynamics of the autotrophs-bacteria interaction in streams – see Figure 1. More specifically, we characterised the effect of enhancing or displacing natural DOC benthic uptake by adding a small flux of labile organic carbon (Lutz et al 2012) and estimating in-stream ecosystem C fluxes under potential stoichiometric constraints (C:N:P) using a flow food web approach (*sensu* Marcarelli et al. 2011).

**Fig 1.**
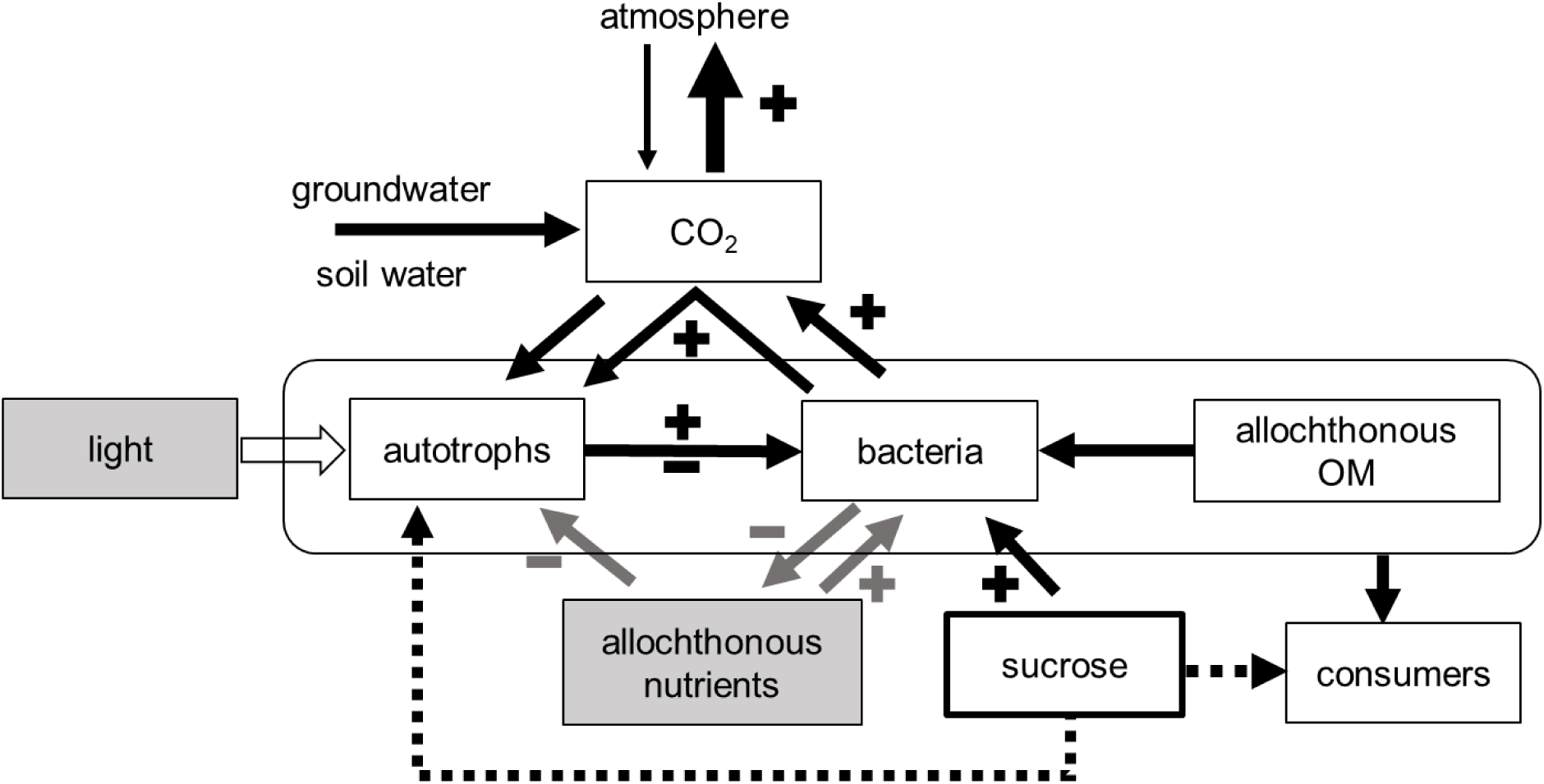
Possible mutual benefits of algae and bacteria and expected changes (+ and − symbols) in carbon and nutrient (N, P) fluxes due to sucrose addition. If nutrients are limiting, then C reciprocal subsidies between bacteria and autotrophs may be weakened by sucrose addition, i.e. there could be a shift between mutualism and competition. With enough nutrients the C microbial loop (to autotrophs) may be strengthened via an increase in CO_2_ from bacterial respiration, i.e. positive feedback loop.

The addition of labile organic carbon (sucrose) may reduce the mineralisation of N and P and increase allochthonous nutrient demand by bacteria, reducing nutrient availability for autotrophs. If nutrients are limiting, then C reciprocal subsidies between bacteria and autotrophs may be weakened by sucrose addition, i.e. there could be a shift between mutualism and competition. With enough nutrients the C microbial loop (to autotrophs) may be strengthened via an increase in CO_2_ from bacterial respiration, i.e. positive feedback loop. Allochthonous nutrients may also come from allochthonous OM so bacteria may be under less nutrient constraints than algae, although the nutrient content of DOC (C:N:P=3200:103:1, Stutter et al. 2013) is generally much poorer than benthic algae (C:N:P=158:18:1, Kahlert 1998). The metabolic balance is likely to shift towards heterotrophy. A large amount of sucrose is expected to be respired, so CO_2_ emissions are predicted to increase. Consumers and the trophic transfer efficiency should benefit from an increase in bacterial production. The design also allows to quantify the potential of priming of allochthonous OM by sucrose via bacterial activity (Kunc et al. 1976, Hotchkiss et al. 2014).

We combined whole-ecosystem stable isotope probing (using sucrose from sugar cane) and Bayesian mixing model to characterise the carbon links (sources to mixtures). We converted the relative carbon fluxes from the different sources, into carbon fluxes by estimating the production of the mixtures, after taking into account assimilation efficiencies (e.g. carbon use efficiency, bacterial growth efficiency). This approach relies on the integration of stream metabolism, nutrient cycling, ecological stoichiometry, stable isotope probing of the food web and production estimates (see Welti et al. 2017). In this study, we focus on the basal part of the food web, and notably on the elusive reciprocal C subsidies between autotrophs and bacteria.

## Methods

### Study area

We studied two heather moorland catchments with soils rich in organic carbon, within the Glensaugh research station of the James Hutton Institute in north-east Scotland (Long 2° 33’ W, Lat 57° 55’ N) – Fig. 2. The streams were about 0.8-1.0 m wide in the studied sections and their channels significantly undercut the banks by 30-46% of stream width. Brown trout (*Salmo trutta fario*, Salmonidae) was present in both streams. The management of the land includes regular heather burning (10-12% of surface area yearly target) for hill farming: mixed grazing of sheep and cattle. For further information see Demars (2018).

**Fig 2.**
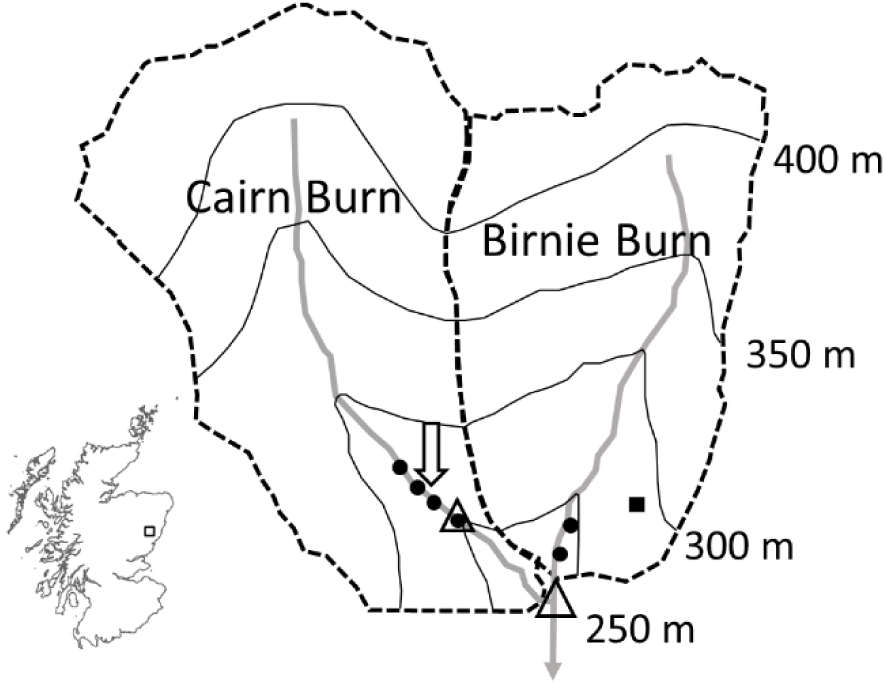
Paired stream experiment at Glensaugh research station. Birnie Burn is the control stream and Cairn Burn the manipulated stream with DOC addition indicated by the arrow. The Cairn Burn also has a control upstream of the treatment reach. The symbols refer to flumes (open triangles), dissolved oxygen stations (filled circles) and soil moisture instrumentation (filled square). The 50 m elevation contour lines are indicated. The catchment area is 0.99 km^2^ (0.90 km^2^ at the flume) for Cairn Burn and 0.76 km^2^ for Birnie Burn. Inset shows the location of Glensaugh in Scotland.

### The control stream

The control stream (Birnie Burn) is part of the long-term monitoring of the UK Environmental Change Network (ECN, http://data.ecn.ac.uk/). There is a monitoring of soil temperature and moisture on the hillslope of the Birnie Burn at 275 m elevation (Cooper et al. 2007). Volumetric soil moisture content is recorded every 30 minutes at 10 and 45 cm depth, corresponding respectively to the base of the O (organic layer) and B (subsoil) horizons of the humus iron podzol present. The stream is equipped with a flume for continuous monitoring of discharge (catchment area 0.76 km^2^) and dip water samples are collected weekly. This stream showed substantial increase in annual flow-weighted mean concentrations of stream water DOC (+0.28 mg C L^−1^ year^−1^ during 1994-2007, Stutter et al. 2011). Water samples were analysed for DOC, pH, nutrients (N, P) and major ions. – see Cooper et al. 2007, Stutter et al. 2012 and Demars 2018 for further details.

### The paired stream

The control stream was paired with a neighbouring stream (Cairn Burn) in 2005, also part of a long-term monitoring scheme with samples collected every week or two for stream water quality. In the late 1970s and early 80s two areas covering 33 ha were improved (reseeded, limed and fertilized) as part of sheep grazing experiments (Hill Farming Research Organisation 1983). The added facilities at the Cairn Burn (catchment area 0.9 km^2^) include a calibrated flume, water level, water electric conductivity, water and air temperature and barometric pressure. Data were recorded every 5 minutes (Campbell Scientific CR10x datalogger). Data logger, battery and barometric pressure were housed in a weather resistant enclosure. Photosynthetic active radiations (PAR) were also recorded in air, one metre above ground, at the same time intervals (LICOR, Lincoln, NE, USA). For more information, see Demars (2018).

### Terrestrial DOC: main source of organic carbon

DOC was the dominant flux of organic carbon (98%) under stable flows with average concentrations of 9.3±1.7 mg C L^−1^ in the two studied streams (Demars 2018). This DOC was of terrestrial origin as shown by δ^13^C analyses of the natural DOC against terrestrial and aquatic plant material (Stutter et al. 2013). The median molar C:N:P stoichiometry of the DOC was 3201:103:1, with values ranging between 978:38:1 to 12013:282:1 (Stutter et al. 2013). Chlorophyll a concentration in the water column was extremely low (< 1 µg L^−1^, Stutter et al. 2013). The pool of particulate organic carbon in the sediment is very small (Demars 2018), and coarse particulate organic carbon was less than 10 g C m^−2^ (determined from Surber sampling of invertebrates, see below).

### DOC addition

A carboy was refilled every two days with 6 kg of sucrose dissolved in over 60 L of stream water filtered through muslin square in a large funnel. The carboy was set as a Mariotte bottle to ensure a constant dripping rate of 22 mL min^−1^ lasting 48 hours (15 mg C s^−1^). The ventilation tube was netted at the top to avoid insect contamination. The dripping rate was kept constant over the 22 days of sucrose addition (23 August - 14 September) and was initially set to increase stream DOC concentration by about 0.5 mg C L^−1^ at 30 L s^−1^. Sample were collected in washed bottles and filtered on-site with pre-washed filters (0.45 μm Millipore PVDF membrane filter). DOC was determined within 48 hours of collection with a Skalar San++ continuous flow analyzer (Breda, The Netherland), using potassium hydrogen phtalate as standards and sodium benzoate for quality controls. The detection limit was 0.1 mg C L^−1^.

### Nutrient cycling studies

Nutrient cycling studies were run in the control and manipulated streams before and during sucrose addition. Nutrient cycling rates were derived from continuous *in-situ* nutrient addition experiments where a conservative tracer is also included (Stream Solute Workshop 1990, Demars 2008). This method tends to overestimate the nutrient uptake length *Sw*, average distance travelled by a nutrient molecule in the water column before river bed uptake. This is due to the addition of nutrient compared to isotopic tracer studies (e.g. Mulholland et al. 2002). However, preliminary tests in the Cairn Burn showed that the bias can be kept small (10-15%) with small nutrient additions (Demars 2008). Nitrate (as KNO_3_) and phosphate (as KH_2_PO_4_) were continuously added together with NaCl as conservative tracer (*cf* Schade et al. 2011). When the plateau phase was reached, water samples were collected at about 10 m interval along the reach and filtered on site (pre-washed 0.45 μm Millipore PVDF membrane filter) (see Demars 2008). The samples were kept cool at 4-10°C. The nutrients (PO_4_ and NO_3_) were determined within 48 hours by colorimetry using a Skalar San++ continuous flow analyzer (Breda, The Netherland) and chloride by ion chromatography (Dionex DX600, Sunnyvale, California). The limits of detections were 0.001 for NO_3_ and PO_4_ and 0.003 mg L^−1^ for Cl. In order to provide a more comparable indicator of nutrient cycling for different hydrological conditions, the uptake velocity *v_f_* was also calculated as follows: *v_f_* = *uz*/*Sw*, with *u* average water velocity and *z* average depth. Short uptake lengths and fast uptake velocities indicate fast cycling rates (high exchange rates between water and benthos).

### Whole-stream metabolism

Whole-stream metabolism was estimated by the open channel two-station diel oxygen method of Odum (1956) modified by Demars et al. 2011b, 2015, 2017 and Demars (2018). Many tracer studies (using NaCl and propane) were carried out as detailed in Demars et al. (2011b) to estimate lateral inflows, mean travel time and reaeration coefficient as a function of discharge within the range of stable flows (up to 32 L s^−1^). The relationships with discharge were very strong (Fig 3, Demars 2018) allowing accurate parameterisation of metabolism calculations under varying flow conditions as in Roberts et al. (2007) and Beaulieu et al. (2013). The high oxygen reaeration coefficient of those streams (0.05-0.24 min^−1^) required very accurate dissolved O_2_ data. Oxygen concentrations were measured with optic sensors fitted on multiparameter sondes TROLL9500 Professional (In-Situ Inc., Ft Collins, CO, USA) and Universal Controller Sc100 (Hach Lange GMBF), the latter powered with two 12 V DC (75mA) car batteries per sensor kept charged with two 20 W solar panels (SP20 Campbell Scientific). The sensors were calibrated to within 1% dissolved oxygen saturation. Four sondes were deployed in the Cairn Burn at 0, 84, 138, 212 m upstream of the flume to include an extra control reach (138-212 m) upstream of the manipulated reach (0-84 m). Another two sondes were set in the control stream Birnie Burn at 88 m interval (60-148 m upstream of the ECN flume). The distances between oxygen stations corresponded to 80-90% of the oxygen sensor footprints (3*u*/*k*_2_), with *u*/*k*_2_ entirely independent of discharge (R^2^=0.0005), which allowed the manipulated reach to be independent of the control reach. The DOC injection point was 28 m upstream of the top station of the manipulated reach, and this distance corresponded to 69% of the oxygen sensor footprint of the top station. All sondes were deployed from May to October 2007, logging at 5 min time step interval.

**Fig 3.**
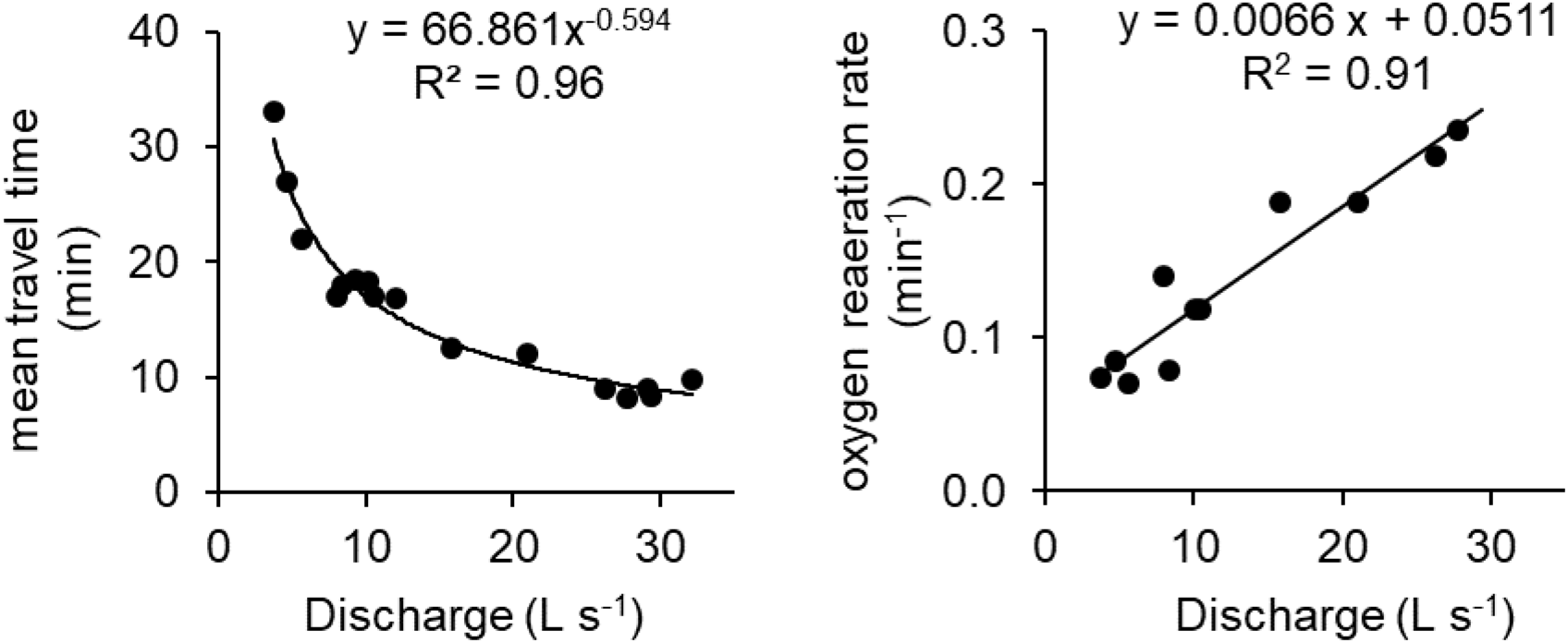
Discharge was an excellent predictor of mean travel time and oxygen reaeration measured from NaCl and propane tracer studies (data from Cairn treatment reach, data of the other two reaches were presented in Demars 2018).

The net metabolism was only calculated for stable flow conditions (3-32 L s^−1^), as follows (Demars 2018):

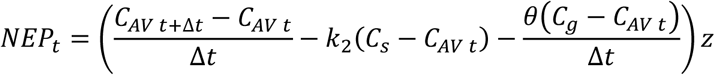

with *NEP_t_* net ecosystem production at time *t*(g O_2_ m^−2^ min^−1^), *C_AV_* average dissolved oxygen (g O_2_ m^−3^) of the two stations at time *t+Δt* and *t*(min), *Δt* time interval (min), *k*_2_ oxygen exchange coefficient (min^−1^), *C_s_* saturated oxygen concentration (g O_2_ m^−3^), *θ* the proportion of lateral inflows, *z* average stream depth (m), *C_g_* oxygen concentration in lateral inflows (g O_2_ m^−3^). The latter was calculated as follows (from baseflow analysis of stream hydrographs):

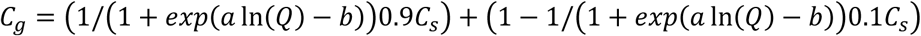

with *Q* discharge, *a* and *b* constants, permitting to correct for baseflow (first term of the equation) and soil water (second term of the equation) lateral inflows, see Demars (2018). The proportion (± se) of total lateral inflows relative to discharge (Q_g_/Q) was 10.7±0.6%, 6.6±0.5%, and 2.3±0.4% for the Birnie Burn control, Cairn Burn control and treated reach, respectively, independently of discharge in the range 3.8-32.5 L s^−1^ (stable flows).

All calculations were run in Excel using a preformatted spreadsheet (Demars 2018). The overall uncertainties in daily stream metabolism, including cross-calibration errors, individual parameter uncertainties, spatial heterogeneity (through the average of diel O_2_ curves) and correction for lateral inflows, were propagated through all the calculations using Monte Carlo simulations (Demars 2018). The corrections for lateral inflows amounted to about 6% of ER for the treated reach (Cairn Burn), 19% and 16% in the control reaches, Cairn Burn and Birnie Burn, respectively.

### Identification of carbon sources and pathways

#### Macrophytes

The percentage cover of bryophytes and filamentous green algae was measured with a ruler across transects taken every two metres along the stream reaches. Young shoots of filamentous green algae and bryophytes were collected by hand along both studied reaches before and after sucrose addition. All samples were freeze dried and milled prior to analyses for C, N, δ^13^C and δ^15^N. The main source of inorganic carbon for primary producers was assumed to be CO_2_ because of the low alkalinity (remaining below 0.5 meq HCO_3_ L^−1^ under low flows). The fractionation factor for CO_2_ assimilation into macrophyte tissue is known to vary with pCO_2_ and growth rate, and was set at −25.5±3.5‰ within the range of Rubisco forms IA and IB in the absence of carbon concentrating mechanism and transport limitation (22–29‰, e.g. Raven et al. 1994, McNevin et al. 2007, Boller et al. 2015). Net primary production was assumed to be driven by filamentous green algae and biofilm autotrophs, and bryophytes were not used to calculate the flow food webs.

#### Inorganic carbon

The δ^13^C of dissolved CO_2_ is known to be variable (Findlay 2004, Billett and Garnett 2010, Billett et al. 2012). The δ^13^C of dissolved CO_2_ reflects both the signature of terrestrial C from groundwater and soil water inflows as well as in-stream processes (biotic respiration) and CO_2_ exchange with the atmosphere (δ^13^CO_2_ about −8‰, e.g. Billett et al. 2012). In the absence of direct measurements, we considered estimating δ^13^C of dissolved CO_2_ from available data of δ^13^C of dissolved inorganic carbon (DIC) in the Brocky Burn, a small stream in the adjacent catchment (Waldron et al. 2007). The equilibrium method used to derive the δ^13^C of CO_2_ from the δ^13^C of DIC (Zhang et al. 1995) was not applicable because the time to reach the carbonate equilibrium (20-200 s) approached the average time spent by a CO_2_ molecule in the stream before emission in the atmosphere (300-1000 s, calculated from travel time and reaeration coefficient of CO_2_, see below). The alternative evasion method (Zhang and Quay 1997) did not seem more appropriate (Billett and Garnett 2010).

We suggest another approach: δ^13^C of dissolved CO_2_ may be estimated indirectly under low flows using the fractionation factor of Rubisco −25.5±3.5‰ and δ^13^C of bryophytes (strict CO_2_ user and no CO_2_ transport limitation). In our study, the average δ^13^C of bryophytes was −36.4‰, and assuming the above fractionation coefficient of −25.5‰, Glensaugh δ^13^C of dissolved CO_2_ would be −10.9‰. This is similar to the δ^13^C of DIC (6-14‰ under low flows, Waldron et al. 2007) and δ^13^C of bryophytes (−33.3‰, Palmer et al. 2001) reported for the Brocky Burn. The proof of concept comes from another Scottish stream (Dighty Burn), where Raven et al. (1994) reported the δ^13^C of dissolved CO_2_ as −14.7‰ and bryophytes as −39.2‰, suggesting a fractionation coefficient of −24.5‰ by difference. In our calculations under low flow conditions we therefore assumed δ^13^C of dissolved CO_2_ as −11±3‰.

To quantify the reciprocal subsidies between autotrophs and bacteria, it remained to decompose the overall stable isotope signature of stream dissolved CO_2_ into the allochthonous (groundwater, soil water and atmospheric exchange) and autochthonous (respiration by heterotrophs and autotrophs) sources. The allochthonous signature, *δ^13^C*_CO2-*aallochthonous*_, can be deduced from rearranging:

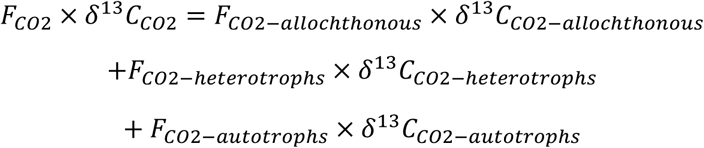

where *F_CO2_* represents CO_2_ fluxes (g C m^−2^ day^−1^, with all fluxes expressed as positive values) and *δ^13^C* the isotope signature (‰) of the different sources of CO_2_. We averaged the δ^13^C of the autotrophs (filamentous green algae and biofilm primary producers). The *δ^13^C*_*CO2-allochthonous*_ was only calculated for the control stream under low stable flows and assumed to apply to both streams. Uncertainties were propagated in quadrature using standard deviation *δx* for sums, and relative uncertainties *δx*/*x* for the division.

#### Sucrose

The δ^13^C of sucrose from sugar cane is similar to that of dissolved CO_2_ (−12±1‰, Jahren et al. 2006, Augspurger et al. 2008, Kankaala et al. 2010, de Castro et al. 2016), but sucrose uptake by autotrophs was assumed to be without isotopic discrimination (Wright and Hobbie 1966). The proportion of carbon derived from added sucrose (F_S_) in resources and consumers was calculated from their δ^13^C in the control (C) and treatment (T) reaches, before (B) and after (A) sucrose addition as follows:

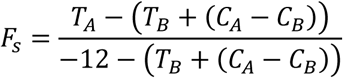

with all uncertainties propagated in quadrature using standard deviation *δx* for sums, and relative uncertainties *δx*/*x* for the division. The standard error of the mean was calculated as 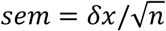 with *n* average number of samples in *C_B_*, *C_A_*, *T_B_*, *T_A_*.

#### Allochthonous organic carbon

The δ^13^C of the DOC (average ±SD) was available from a previous study from the same catchment and showed it was of terrestrial origin, i.e. not autochthonous (δ^13^C = −28.5±0.3 ‰, Stutter et al. 2013, see above). Coarse particulate organic matter (CPOM) was also collected by hand along both studied reaches before and after sucrose addition. Since there was hardly any difference in δ^13^C between DOC and CPOM (δ^13^C = −27.4±0.7 ‰, Table S1), we used the δ^13^C of CPOM determined in this study as the signature for allochthonous organic carbon.

#### Periphyton

Periphyton (or biofilm) samples represent a mixture of primary producers (algae and cyanobacteria), bacteria and fine particulate organic matter. The samples were collected before and at the end of the sucrose addition from the flumes and stones with a toothbrush, funnel and bottle. All samples were freeze dried and milled.

We also placed six pairs (with/without Vaseline) of unglazed ceramic tiles (10 × 10 cm) fixed on bricks and deployed along the studied reaches in both streams three weeks before the start of the manipulation. Vaseline was applied around half the tiles to prevent grazing by invertebrates. After three weeks, there was hardly any growth on the tiles, and so the tiles were left in the stream until the end of the manipulation. One brick in the control stream was lost. At the end of the experiment, the tiles were frozen at −20°C, later freeze dried and the biofilm was scraped with a razor blade. Since there was little biomass per tile (about 1 g C m^−2^), the biofilm was pooled by stream and grazer treatments (leaving two samples per stream). Very little grazing activity was observed on the tiles during the six weeks and unsurprisingly no difference in biofilm dry mass emerged due to grazer exclusion (paired t-tests on ln transformed mass; Birnie, t_4_, p=0.13; Cairn, t_5_, p=0.26).

Phospholipid fatty acids (PLFAs) were extracted from the biofilm samples from the tiles and derivatised to their methyl esters (FAMES) following the procedure of Frostegård et al. (1993) using the modified extraction method of Bligh and Dyer (1959), as detailed in Certini et al. (2004). Quantification and δ13C values of the PLFAs were both determined by Gas Chromatography-Combustion-Isotope Ratio Mass Spectrometry (GC-C-IRMS) as described by Main et al. (2015), and averaged for each stream (δ^13^C, Table S2). Only PLFAs up to 19 carbon chain length were determined (excluding some long-chain essential polyunsaturated fatty acids, e.g. Muller-Navarra et al. 2000, Gladyshev et al. 2011).

#### Bacterial carbon

The bacterial PLFAs were identified as those most affected by sucrose addition in the treated reach and information derived from the literature (supplementary information). This also allowed to determine the δ^13^C of bacteria in the control reach. We used a fractionation factor of −3‰ for the δ^13^C of bacterial fatty acids relative to bulk tissue samples (Hayes 2001, Gladyshev et al. 2014). This is within the range of observed values in other studies (Boschker and Middelburg 2002, Bec et al. 2011). We used the same fractionation factor of −25.5±3.5‰ for the assimilation of CO_2_ coming from bacterial respiration into green algal tissue. Since we only had comparative data for the treatment period, the fraction of sucrose within PLFAs was calculated as follows:

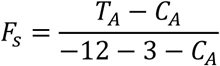

#### Biofilm autotroph carbon

The carbon from cyanobacteria and algae were identified from a specific PLFA (α-linolenic acid, Risse-Buhl et al. 2012) using the same fractionation factor as above between PLFA and bulk tissue (Table S2).

#### Macroinvertebrate consumers

Macroinvertebrate densities were estimated from twelve to thirteen Surber samples (20 × 20 cm, mesh size 200 µm) collected randomly along the reaches (total 51 samples). The samples were stored in 70% alcohol, sorted and identified. Macroinvertebrate for CN and stable isotope studies were collected by kick sampling and hand net. The animals were sorted and identified live within a day into Petri dishes and after allowing time for gut evacuation, placed in Eppendorf tubes and freeze dried. The average biomass of macroinvertebrate taxa was assessed by weighing the freeze-dried mass of all individuals within a tube divided by the number of animals. The whole macroinvertebrates were then crushed before insertion into a tin capsule.

#### Carbon isotopic turnover

Bacteria and algae were likely to be fully labelled within a week, and three weeks of sucrose addition was thought to be sufficient to label invertebrates, albeit not fully for all consumer species (especially predators), with carbon turnover ranging from about 10-35 days (Le Cren and Lowe-McConnell 1980, Hall and Meyer 1998, Collins et al. 2016 and references therein), and a time lag to reach equilibrium at each trophic level (e.g. the time for bacteria and algae to reach full equilibrium, likely to be less than a week, will delay the time grazers may reach their full equilibrium). The proportion of tissue turnover over 14 and 21 days was estimated from the consumer individual body mass *M*(g) of the taxa (assuming freeze-dried mass = 0.2 fresh mass, Waters 1977 cited in Wetzel 2001, p. 718) with tissue isotopic turnover rate λ (day^−1^) derived from the isotopic half-life study of Vander Zanden et al. (2015), general equation for invertebrates:

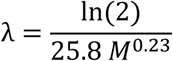

and from the remaining individual body mass *M_t_* at time *t*:

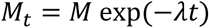

This provided an indication to what extent the isotopic signature of consumers may have reached an equilibrium. We used the proportion of macroinvertebrate tissue turnover over 21 days to provide the fraction of sucrose at equilibrium for individual taxa and grouped by functional feeding groups following Demars et al. (2012).

#### Analytical methods

The total carbon and total nitrogen concentrations and the δ^13^C and δ^15^N natural abundance isotope ratios of the milled samples were determined using a Flash EA 1112 Series Elemental Analyser connected via a Conflo III to a Delta^Plus^ XP isotope ratio mass spectrometer (all Thermo Finnigan, Bremen, Germany). The isotope ratios were traceable to reference materials USGS40 and USGS41 (both L-glutamic acid); certified both for δ^13^C (‰_VPDB_) and δ^15^N (‰air N2). The carbon and nitrogen contents of the samples were calculated from the area output of the mass spectrometer calibrated against National Institute of Standards and Technology standard reference material 1547 peach leaves which was analysed with every batch of ten samples. Long term precisions for a quality control standard (milled flour) were: total carbon 40.3 ± 0.35 %, δ^13^C −25.4 ± 0.13 ‰, total nitrogen 1.7 ± 0.04 % and ^15^N 0.367 ± 0.0001 atom % (mean ± sd, n = 200). Data processing was performed using Isodat NT software version 2.0 (Thermo Electron, Bremen, Germany) and exported into Excel. Total P was determined after 30 min digestion in 50% nitric acid at 120°C for CPOM, bryophytes, biofilm and green filamentous algae (see Demars and Edwards 2007).

#### Data analyses

Most studies use δ^13^C and δ^15^N to identify the flow path in the food web. Here the BACI experimental design allowed to calculate the proportion of sucrose (Fs) in all parts of the food web and was used as a tracer in addition to δ^13^C to determine the sources of carbon for bacteria and algae in the treatment reach after 21 days of sucrose addition. Thus, the carbon pathways were identified with carbon tracers. End member mixing analyses were used to determine the proportion of C sources and their uncertainties in primary producers and bacteria. We provided the numerical solutions given by a Bayesian stable isotope mixing model SIAR 4.2.2 (Parnell et al. 2010) in R version 3.1.3 (R Core Team 2015). The numerical solutions of SIAR were very similar to the analytical solutions of IsoError 1.04 (Phillips and Gregg 2001), and suggested the results were not biased (*cf* Brett 2014). Note, the relative importance of individual resources to consumers was outside the scope of this study.

### Quantification of carbon fluxes

To assess trophic transfer efficiency through the food web, our production estimates were all standardized to g C m^−2^ day^−1^. Respiration and photosynthesis rates in oxygen were converted to carbon using a respiratory and photosynthetic quotient of 1 (Williams and del Giorgio 2005, but see Berggren et al. 2012).

#### Total CO_2_ emissions

In the absence of direct measurements, the excess partial pressure of CO_2_ (E*p*CO_2_) of the streams was estimated from three measured parameters: pH, alkalinity and temperature (Neal et al. 1998, as applied in Demars et al. 2016 with atmospheric CO_2_=384 ppm, ftp://ftp.cmdl.noaa.gov/ccg/co2/trends/co2_annmean_mlo.txt). Our E*p*CO_2_ estimates have high uncertainties (±50%, Demars 2018). E*p*CO_2_ is the concentration of free CO_2_ in the stream water (*C_t_* at time *t*) relative to the atmospheric equilibrium free CO_2_ concentration (*C_SAT_*):

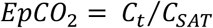

*C_SAT_* was calculated from published CO_2_ solubility in pure water at equilibrium with atmospheric CO_2_ in the temperature range 0-90°C (Carroll et al. 1991) and Henry’s law (Stumm and Morgan 1981, Butler 1982). *C_t_* was calculated as EpCO_2_ × *C_SAT_*. The flux of CO_2_ (*F_CO_2__*, g C m^−2^ day^−1^) at the interface between water and the atmosphere was calculated as for O_2_ following Young and Huryn (1998):

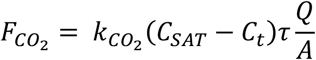

with *k_co_2__* reaeration coefficient of CO_2_ (day^−1^), *C_SAT_* and *C_t_* (mg C L^−1^ or g C m^−3^), *τ* mean travel time of the stream reach (day), *Q* average water discharge (m^3^ day^−1^), *A* surface water area of the stream reach (m^2^). The reaeration coefficients between CO_2_ and O_2_ can be simply related as follows (Demars et al. 2015, Demars 2018):

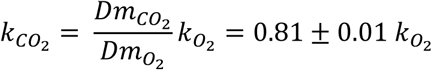

based on the molecular diffusivity (*Dm*) of CO_2_ and O_2_ measured at three different temperatures within the same study (Davidson and Cullen 1957).

The flux of CO_2_ was then related to discharge within the range of low stable flows for which stream metabolism was processed (Cairn n=47, R^2^=0.81; Birnie n=65, R^2^=0.77) to provide daily estimates. For more details, see Demars 2018.

#### Biotic CO_2_ emissions

These were simply calculated as the net ecosystem production (NEP), gross primary production (GPP) plus ecosystem respiration (ER, a negative flux) expressed in g C m^−2^ day^−1^. Bacterial respiration of DOC was calculated as heterotrophic respiration (HR, a negative flux) from:

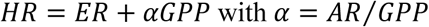

with AR, autotrophic respiration and ER, ecosystem respiration, both negative fluxes (oxygen consumption) and GPP, positive flux (producing oxygen). We partitioned ER into auto and heterotrophic respiration with α=0.5 (see Demars et al. 2015) and calculated uncertainties using α=0.2 and α=0.8. Bacterial respiration of the added sucrose was calculated as the difference in heterotrophic respiration between the treatment and a control reach during sucrose addition, after standardising for site differences using the control period.

#### Allochthonous organic matter

The overall flux at the outlet of both streams was calculated as discharge times DOC concentration using the weekly data from the long-term ECN monitoring collected in 2007-2008 under low stable flows. The DOC flux was then related to discharge (Cairn n=74, R^2^=0.85; Birnie n=100, R^2^=0.83), to provide an estimate for the days for which DOC was not measured (within the same range of flows).

The organic carbon uptake length (*Sw_OC_*, in m) and mineralisation velocity (v_f-OC_, in m day^−1^) were calculated as in previous studies (Newbold et al. 1982, Hall et al. 2016), here neglecting POC (see above):

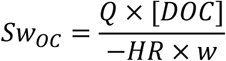

with [DOC] dissolved organic carbon concentration (g C m^−3^ day^−1^), Q discharge (m^3^), HR heterotrophic respiration (a negative flux expressed in g C m^−2^ day^−1^) and w width (m), and

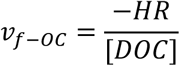

#### Light

We derived a conversion factor for photosynthetically active radiation (PAR; 1 mol photon m^−2^ day^−1^ = 6.13 g C m^−2^ day^−1^) by relating the ratio of total quanta to total energy within the PAR spectrum (2.5 × 10^21^ photon s^−1^ kJ^−1^ = 4.15 × 10^−3^ mol photon kJ^−1^; Morel and Smith 1974) with the reciprocal of the energy content of glucose expressed in carbon units (15.7 kJ g^−1^ glucose = 25.4 × 10^−3^ g C kJ^−1^; Southgate and Durnin 1970).

#### Sucrose

The flux of added sucrose over the treatment reach was determined at the top of the reach (84 m upstream of the flume of Cairn Burn) using the average observed increase in DOC concentrations (28/08, 5/09, 11/09/2007) multiplied by discharge (Fig S1).

#### Whole-stream production

We calculated the net ecosystem primary production (NPP) by assuming a 50% carbon use efficiency ε, that is half GPP (reviewed in Demars et al. 2015, 2017) and computed NPP for ε=0.2 and ε=0.8 to provide uncertainties in our estimates. Heterotrophic production (HP, g C m^−2^ day^−1^) was calculated from heterotrophic growth efficiency HGE and heterotrophic respiration (HR, negative flux expressed in g C m^−2^ day^−1^) as follows:

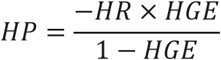

Heterotrophic production was estimated for low (5%) and moderate (20%) bacterial growth efficiencies (e.g. Berggren et al. 2009, Fasching et al. 2014, Berggren and del Giorgio 2015).

#### Macroinvertebrate consumers

Secondary production was estimated from the samples collected at the end of the treatment period from the observed standing biomass (mg C m^−2^) of individual taxa and macroinvertebrate daily growth rate (day^−1^). The standing biomass was determined from the density (individuals m^−2^) and average individual biomass (mg C). The growth rate (*G*) was determined from:

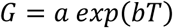

with *T* average water temperature (10.5°C), *a* and *b* taxon specific constants derived from a global compilation of published data (Golubkov 2000, Gladyshev et al. 2016 – similarly to Morin and Dumont 1994).

### Ecosystem efficiencies

With all fluxes expressed in g C m^−2^ day^−1^, we calculated the light use efficiency (net primary production/photosynthetic radiations), organic carbon use efficiency (bacterial production/DOC supply), resource use efficiency (consumers/ [net primary production + bacterial production]), and the proportion of biotic CO_2_ emissions (net ecosystem production / total CO_2_ emissions).

### Data analyses

We calculated an effect size (i.e. proportional changes) of sucrose addition on our response variables (e.g. nutrient cycling, stoichiometric ratios, metabolic fluxes, ecosystem efficiencies) using the values of the control (C) and treatment (T) reaches, before (B) and after (A) sucrose addition as follows:

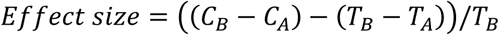

with all uncertainties propagated in quadrature using standard deviation *δx* for sums, and relative uncertainties *δx*/*x* for the division. The standard error (*se*) of the effect size was calculated as 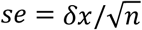 with *n* average number of independent samples or measurements in *C_B_*, *C_A_*, *T_B_*, *T_A_* (although strictly speaking the samples were pseudo-replicated because the random collection was within one plot). Since the overall design was unreplicated we simply interpreted the effect size in relation to the standard error (we do not report p values).

The control period for the metabolic parameters and trophic transfer efficiencies was limited to the two days prior to sucrose addition (21-22 August), so the whole data series was within a single period of low stable flows, i.e. not interrupted by peak flows, producing more comparable results across sites. Daily metabolic parameters were temporally pseudo-replicated (especially during the control period), so it was only possible to report relative uncertainties (*δx*/*x*).

All effect sizes and efficiencies were calculated at the reach scale.

## Results

### DOC addition

We were fortunate to have no rainfall and low stable flows during the whole three weeks of carbon addition. The target concentration was achieved at the top of the studied reach, 28 m downstream of the injection point, where the DOC addition averaged 0.52 mg C L^−1^ (Fig. S1), despite a lower than expected discharge (from 20 L s^−1^ down to 8 L s^−1^ by the end of the addition), due to significant uptake and mineralisation within the 28 m mixing zone (an average 55% loss of sucrose flux from the point of injection to the top of the reach).

### Stream metabolism continuous monitoring

Bryophytes covered 4, 11 and 17% of the river bed in the Birnie control, Cairn control and Cairn treatment reach, respectively. Filamentous green algae (mostly *Microspora* sp., Microsporaceae) percentage cover increased during the period of sucrose addition from 1 to 19%, 11 to 33% and 4 to 37% of channel width in the Birnie control, Cairn control and Cairn treatment reach, respectively.

Gross primary production (GPP) peaked to 7.6 g O_2_ m^−2^ day^−1^ ten / eleven days after the start of sucrose addition in the treatment reach before decreasing sharply down to an average 2.4 g O_2_ m^−2^ day^−1^ during the last four days of the carbon addition, and this despite high photosynthetic active radiations. In contrast, GPP remained relatively constant in the control reaches Birnie Burn (about 1.2 g O_2_ m^−2^ day^−1^) and Cairn Burn (3.2 g O_2_ m^−2^ day^−1^ for the first two weeks declining to 2.1 g O_2_ m^−2^ day^−1^ during the last week g O_2_ m^−2^ day^−1^) – Fig. 4.

**Fig. 4.**
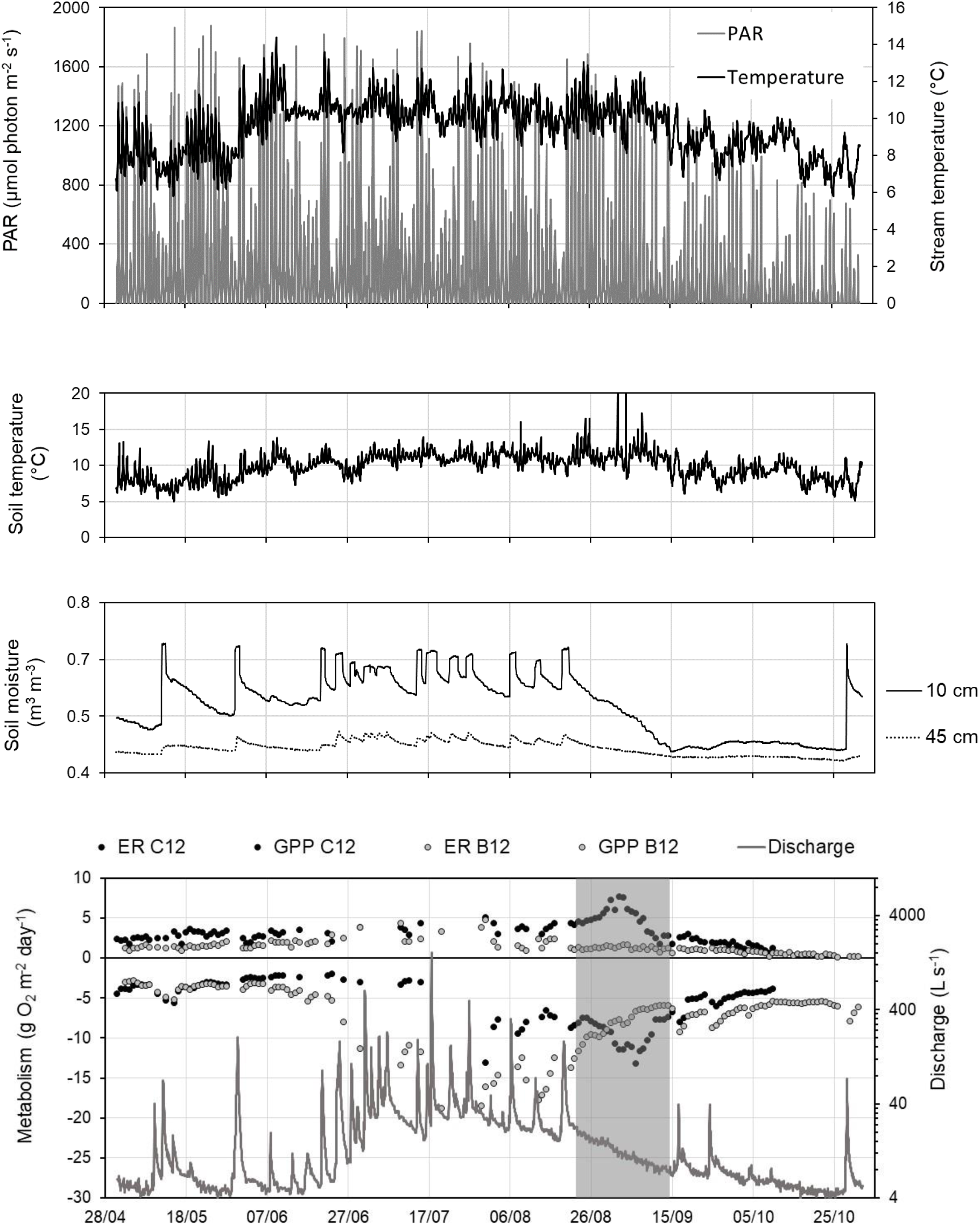
Continuous data monitoring and stream metabolism in the control reach (Birnie Burn, B12) and treatment reach (Cairn Burn, C12). Ecosystem respiration (ER) is negative (consuming O_2_) and gross primary production (GPP) is positive (producing O_2_). The period of DOC (sucrose) addition (23/08-14/09) is indicated by grey shading. The depths for soil moisture correspond to the depths of the organic soil (10 cm) and subsoil (45 cm).

Peaks in ecosystem respiration (ER) down to −20 g O_2_ m^−2^ day^−1^ (Birnie Burn) and −35 g O_2_ m^−2^ day^−1^ (Cairn Burn) were more visible for the controls than the treatment reach, with respiration activity inversely related to soil hydrological connectivity, as recorded by soil moisture continuous monitoring (Fig. 4). There was an increase in ER in the treatment reach at the start of the sucrose addition, despite the continuing loss of hydrological connectivity. More specifically, heterotrophic respiration activity associated to sucrose addition peaked sharply 15 days after the start of the addition, processing up to 59% of the daily sucrose flux (Fig. 5). On average 35±20% of the added sucrose was respired during the addition over just 84 m (or 15 minutes mean travel time). Heterotrophic production ranged between 2% and 10% of the sucrose flux, based on bacterial growth efficiencies of 0.05 and 0.2 respectively.

**Fig 5.**
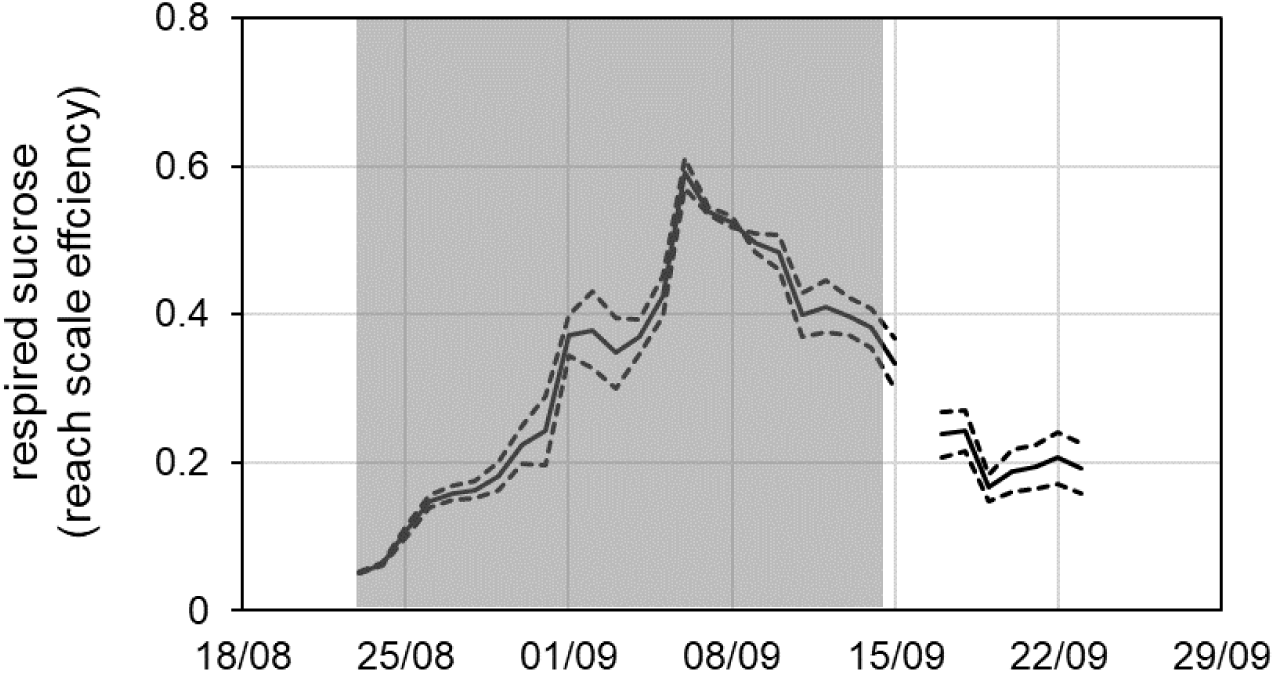
Efficiency of respired sucrose in the 84 m long treatment reach (with mean travel time of 15 minutes) during (shaded area, 23/08-14/09) and shortly after sucrose addition, calculations based on heterotrophic respiration in the treatment reach relative to the Birnie control (same results with Cairn control, not shown). The black line was calculated with an autotrophic respiration (AR) of 0.5×GPP and dashed lines with AR=0.2×GPP and 0.8×GPP (see method). On average 35±20% of the daily flux of sucrose was respired within that reach during the sucrose addition.

### Nutrient cycling studies and stoichiometry

The background concentrations of nitrate and phosphate were 180 and 90 µg N L^−1^ and 2 and 4 µg P L^−1^ in the Birnie control and Cairn treatment reach, respectively. The added geometric mean of N and P were on average 471 µg N L^−1^ and 24 µg P L^−1^. The addition of sucrose had no effect on nitrate and phosphate nutrient uptake length and uptake velocity (Fig. 6, Table S3). The phosphate uptake length was highly related to discharge and became very short, down to 31 m in the treatment reach towards the end of the sucrose addition. Phosphate uptake velocity was about 0.2 mm s^−1^ and an order of magnitude faster than nitrate with uptake lengths in the kilometre range.

**Fig 6.**
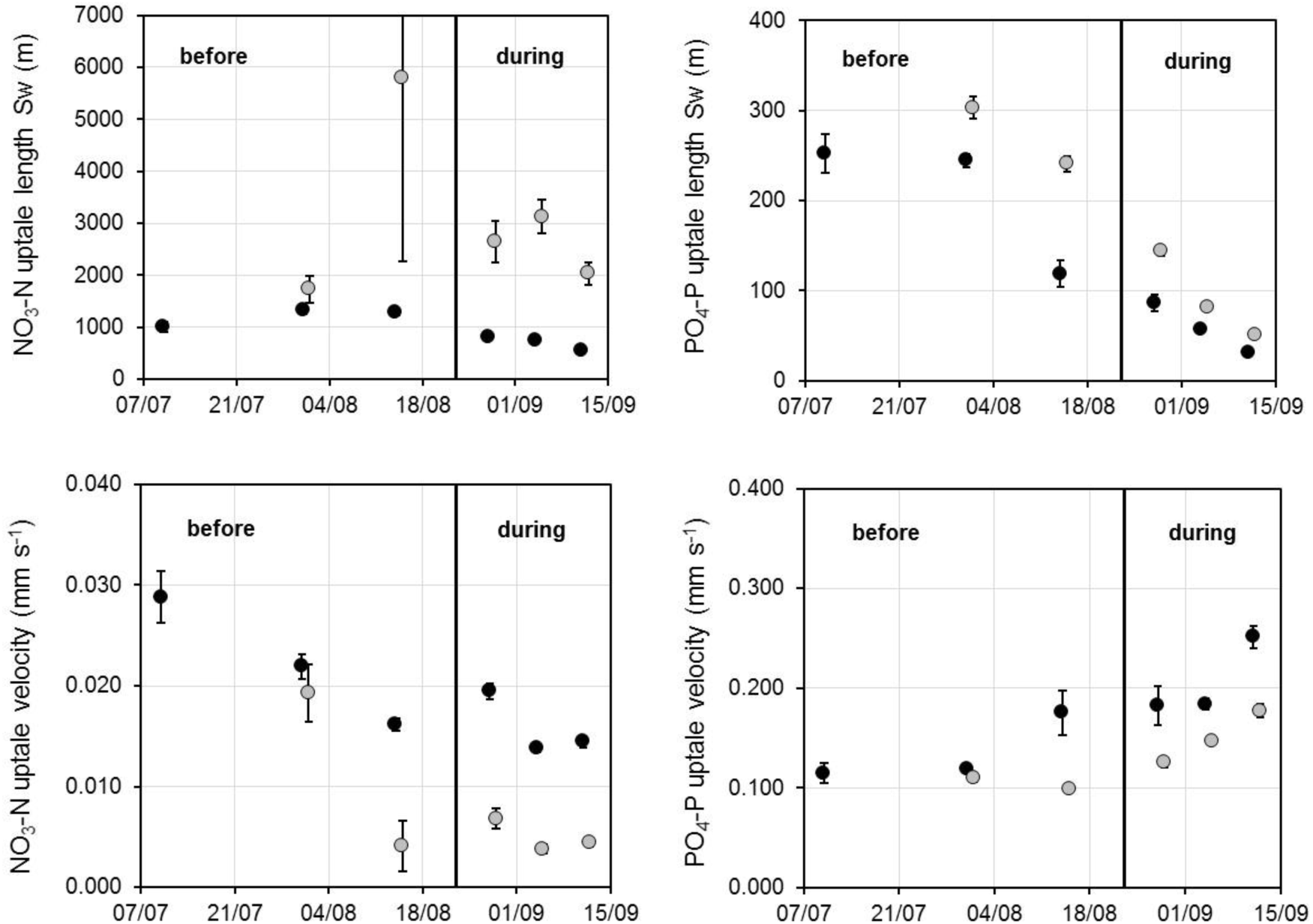
Stream nutrient uptake length (S_w_) and uptake velocity (v_f_) before and during sucrose addition in the control (grey symbols) and treatment (black symbols). Error bars represent sem, with some error bars smaller than the symbols. Note the different magnitudes on the y axes between NO_3_-N and PO_4_-P.

The molar C:N:P stoichiometric ratios of coarse particulate organic matter and bryophytes remained stable throughout the experiment. Sucrose addition exerted strong effects on filamentous green algae and periphyton stoichiometry (Fig. 7, Table S4). While the molar C:N:P stoichiometric ratios decreased in the control stream, they increased sharply in the treatment reach following sucrose addition: from 330:29:1 to 632:49:1 in filamentous green algae and from 262:29:1 to 428:38:1 in periphyton.

**Fig 7.**
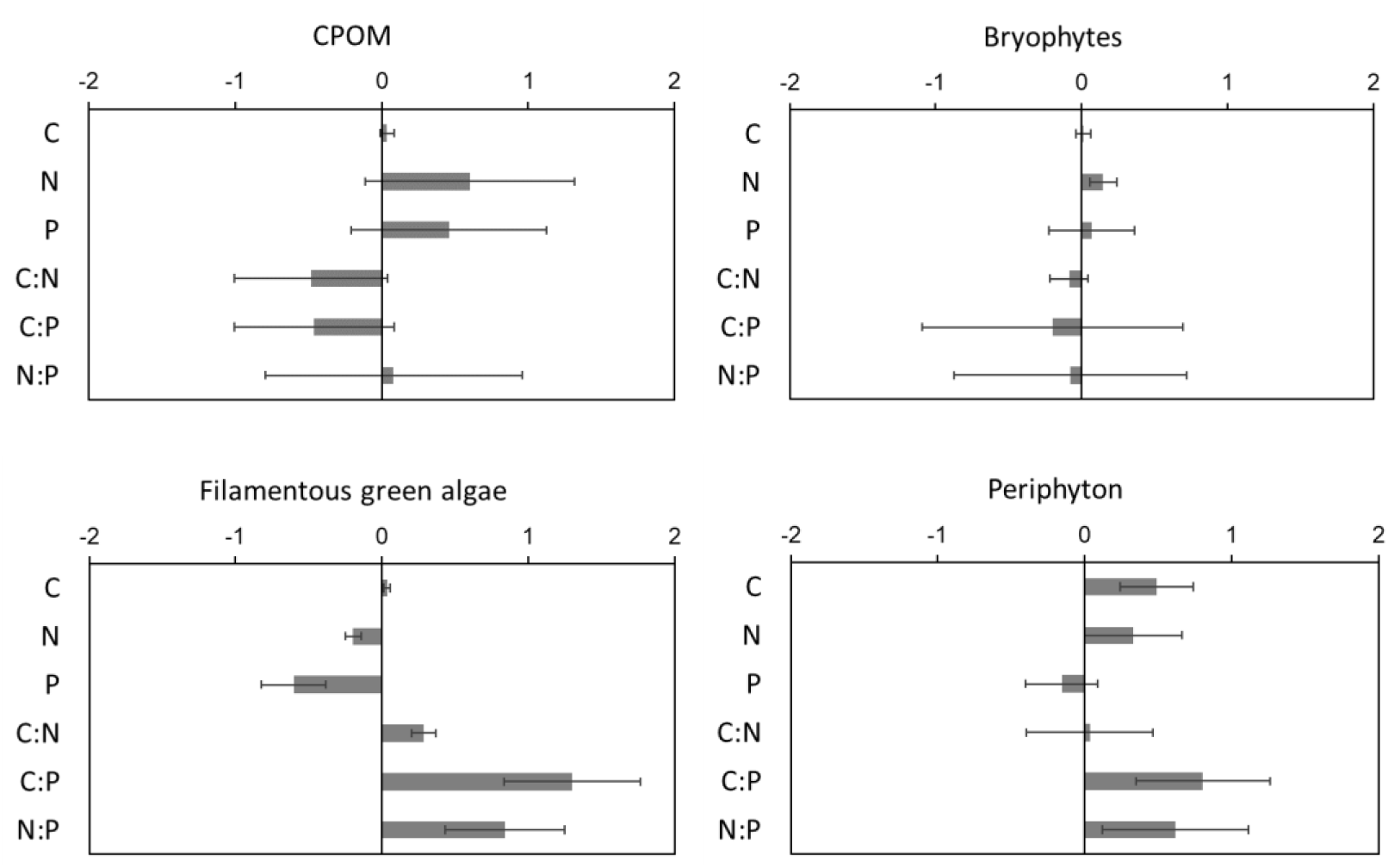
Proportional changes in C, N, P (% w/w) and molar C:N:P stoichiometric ratios in the basal food web resources due to sucrose addition based on the before and after control impact experimental design. Error bars represent sem.

### Fate of added sucrose

The fraction of carbon derived from sucrose (F_S_) among the food web resources filamentous green algae, periphyton and its autotrophs and heterotrophs (bacteria) was relatively high at 24, 23, 36 and 68%, respectively (Fig. 8, Table S1). The average (range) tissue turnover of consumers was 76% (53-92%) to 88% (67-98%) over 14 to 21 days. The proportion of carbon derived from sucrose in macroinvertebrates, after correcting for tissue turnover, varied among taxa but averaged around 23% independently of the functional groups, except for filter feeders with F_S_=64% (Fig. 9, Table S5).

**Fig. 8.**
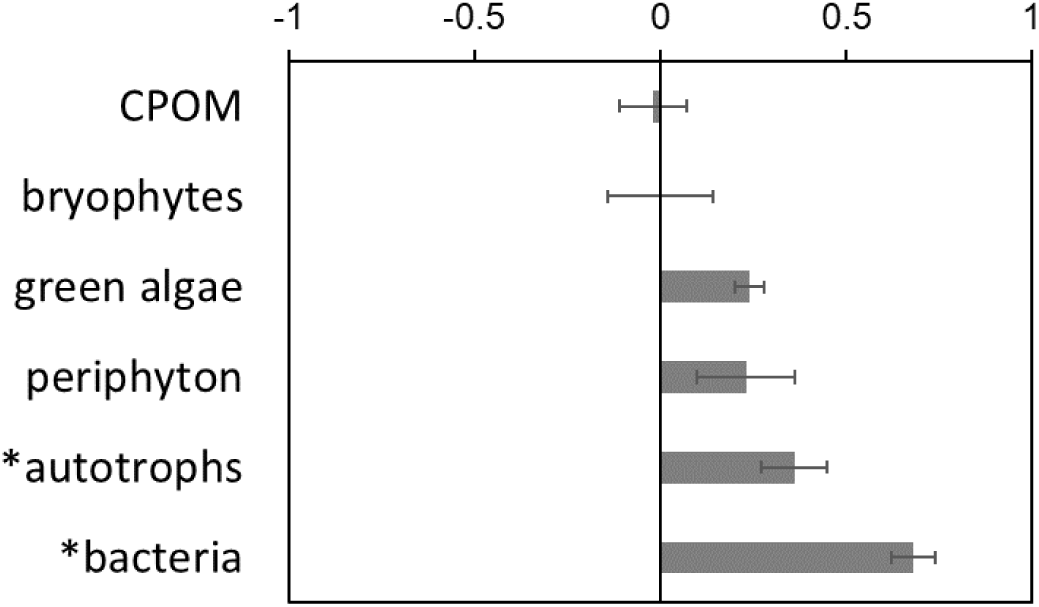
Proportion of carbon derived from added sucrose in food web basal resources based on δ^13^C (see TABLE S1) and PLFA δ^13^C for autotrophs and bacteria in periphyton (indicated by *, See TABLE S2). Error bars represent se. Size effect calculated from BACI design, except for periphyton autotrophs and bacteria (periphyton collected at the end of the experiment in the control and treatment reach, see method).

**Fig. 9.**
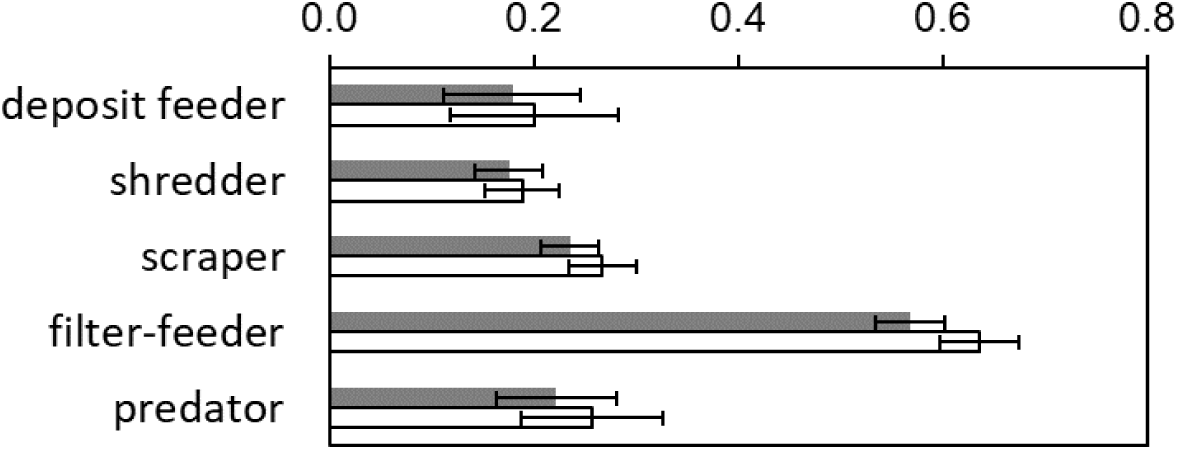
Proportion of carbon derived from added sucrose in macroinvertebrate consumers by functional feeding groups based on the before and after control impact experimental design and associated δ^13^C changes (see Table S5); observed (grey bars) and at equilibrium (open bars, see method); error bars represent the standard error.

### Identification of carbon sources and pathways under low flows (end of experiment)

The autotrophs (filamentous green algae and periphyton autotrophs) derived 54±11% of their CO_2_ from allochthonous sources (groundwater and atmosphere) and 46±11% from bacteria in the control reach. In the treatment reach, autotrophs derived a similar proportion of CO_2_ from allochthonous sources (59±9%), some osmotrophic uptake of sucrose (10±7%), and relatively less from bacteria (31±11%). The bacteria used mostly autotrophic carbon (76±12%) relative to allochthonous organic carbon (24±12%) in the control, but preferred sucrose (51±7%) to autotrophic (34±12) and allochthonous organic matter (15±12%) in the treatment reach – see Fig. S2.

### Quantification of carbon fluxes and efficiencies

The flow food web of the control and treatment reach over the three weeks of sucrose addition were quantified according to our conceptual model (see Fig. 1). Figure 10 illustrates the C fluxes of net primary production, bacterial production and secondary production as well as bacterial CO_2_ flux and overall net ecosystem production (emission of CO_2_ to the atmosphere). Photosynthetic active radiation (light) decreased slightly from 106 to 80 and 118 to 90 g C m^−2^ day^−1^ in the control and treatment respectively. Allochthonous organic matter (DOC) was more than halved from 177 to 85 and 110 to 50 g C m^−2^ day^−1^ in the control and treatment. This was reflected by a general reduction in the organic carbon uptake length 3214 to 2531 m and 4257 to 1886 m in the control and treatment, respectively, independently of the carbon addition. The organic carbon uptake velocity decreased in the control from 0.82 to 0.55 m day^−1^ but increased in the treatment from 0.53 to 0.76 m day^−1^.

**Fig 10.**
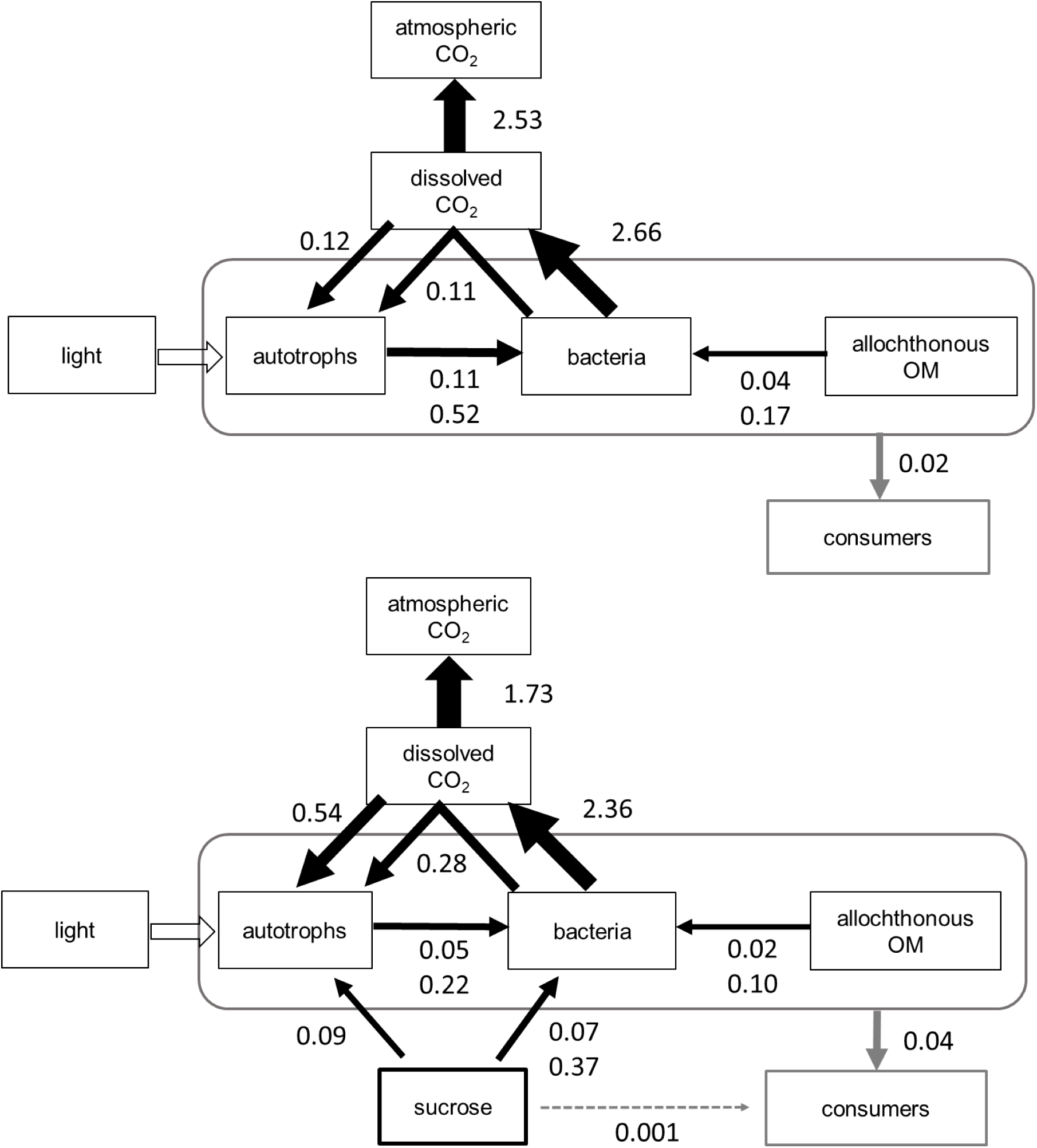
**Flow food webs under low flows when stream water hydrologically disconnected from soils:** in-stream biotic carbon fluxes (g C m^−2^ day^−1^, black and grey arrows) and energy flow (light) in the control (top) and treatment reach (bottom) after three weeks of sucrose addition, based on source partitioning using stable isotopes and production estimates. Autotrophs represent net primary production, bacteria represent heterotrophic production, and consumers represent secondary production. Bacterial respiration and net ecosystem production (biotic CO_2_ emissions) are also represented. Two figures were given for bacterial production based on heterotrophic growth efficiencies (HGE) of 0.05 (low) and 0.2 (moderate). Note: CO_2_ emissions to the atmosphere represented 26% and 12% of total CO_2_ emissions, in the control and treatment respectively with most CO_2_ derived from soil water and groundwater.

Our estimates suggest that a large part of net primary production (0.23, range 0.09-0.38 g C m^−2^ day^−1^) was used for bacterial production (0.15-0.69 g C m^−2^ day^−1^) in the control, and in turn nearly half of the CO_2_ fixed by autotrophs was derived from bacterial CO_2_, although it represented a small fraction of bacterial respiration driving net ecosystem production (biotic CO_2_ emissions). These reciprocal C subsidies between autotrophs and bacteria were not as strong relative to net primary production (0.91, range 0.36-1.46 g C m^−2^ day^−1^) and bacterial production (0.14-0.66 g C m^−2^ day^−1^) in the treatment during sucrose addition. The estimated flux of allochthonous organic matter assimilated by bacteria was similar in the control (0.04-0.17 g C m^−2^ day^−1^) and treatment (0.02-0.10 g C m^−2^ day^−1^). Part of the allochthonous organic matter respired by bacteria was recycled by primary producers and accounted for 11±6% of net primary production in the control.

We also derived, from the BACI design, the effect size of sucrose addition on selected whole-ecosystem metabolic properties and efficiencies (see Fig. 11, Table S6). All estimated ecosystem properties and efficiencies were summarised in Table S7 for Birnie control and Cairn treatment before and after sucrose addition. We present some key highlights below.

**Fig. 11.**
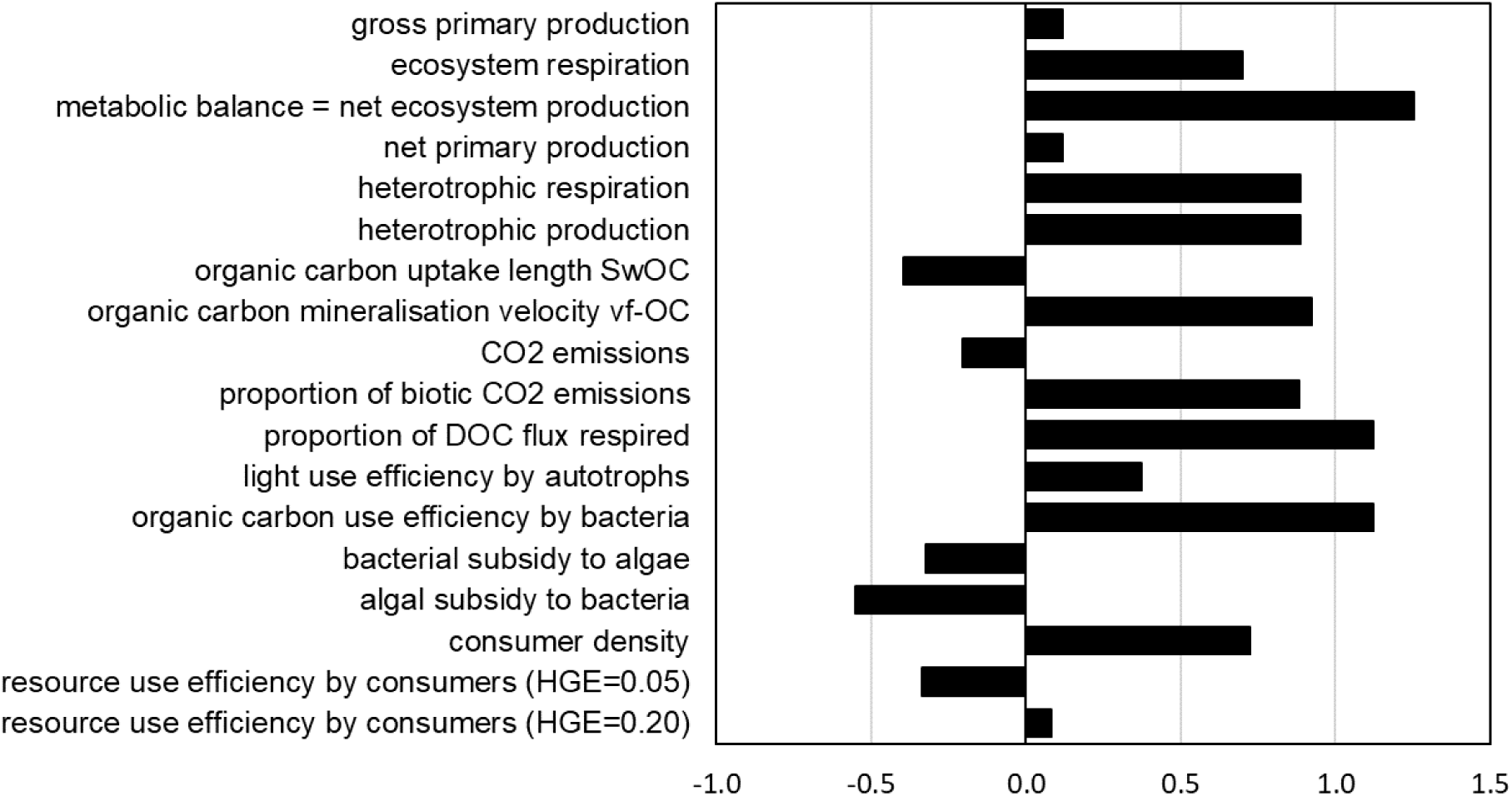
Effect size (1=100%) of sucrose addition on selected ecosystem properties at reach scale based on the before and after control impact experimental design, except for resource use efficiency by consumers and algae-bacteria reciprocal subsidies relying on a simple comparison between the control and the treatment reach. See Table S6 for uncertainties (most very large) and Table S7 for the individual values. The resource use efficiency by consumers was estimated for two heterotrophic growth efficiency (HGE).

While GPP increased marginally (12%), there was a small relative increase in light use efficiency (37%). ER intensified by 70%, and the net ecosystem production (NEP) became relatively more negative by 125%, i.e. 125% relative increase in biotic CO_2_ emissions. Heterotrophic respiration and production increased by 89%, and this was reflected by a shorter (−40%) uptake length (Sw_OC_) and faster mineralisation velocity (92%) of organic carbon. The proportion of DOC flux respired (range 2.2-5.3%) and organic carbon use efficiency by bacteria (range 0.1-0.3%) increased by 112%. While there was a relative decrease (−20%) in total CO_2_ emissions, the proportion of biotic CO_2_ emission increased by 88%. The reciprocal subsidies between autotrophs and bacteria were weaker by 33% (autotrophs to bacteria) and 55% (bacteria to autotrophs) in the treatment relative to the control.

The average consumer biomass per individual was similar between the control (0.20 mg C ind^−1^) and the treatment reach (0.21 mg C ind^−1^) and average production per individual was slightly higher in the treatment reach (6 µg C ind^−1^ day^−1^) than in the control (5 µg C ind^−1^ day^−1^), at the end of the sucrose addition. Consumer density (range 1300-6000 ind m^−2^) increased by 72% due to sucrose addition and consumer production was higher in the treatment (36 mg C m^−2^ day^−1^) than in the control (20 mg C m^−2^ day^−1^) at the end of the experiment. The resource use efficiency (trophic transfer efficiency) by consumers was similar between the control and the treatment reach (2-5%), with a size effect of sucrose addition ranging from −33% to +8% depending on the heterotrophic growth efficiency (0.05 and 0.2, respectively) used to calculate heterotrophic production.

## Discussion

Our experiment showed that a small continuous addition of labile DOC (0.52 mg C L^−1^ as sucrose, 12% of total DOC, Fig. S1) can profoundly alter whole-ecosystem behaviour. The use of a before and after control experiment together with the addition of a deliberate tracer with a distinctive δ^13^C signature allowed not only to trace the fate of the added carbon into the treated reach but also to build the flow food web of the control reach, unravel C reciprocal subsidies between autotrophs and bacteria, and demonstrate the potential for some natural allochthonous organic matter to feed the primary producers via bacterial respiration.

### Reciprocal subsidies between autotrophs and bacteria

The use of autotroph carbon by bacteria has been shown before using a photosystem II inhibitor in biofilm (Neely and Wetzel 1995) or δ^13^C_DIC_ additions (e.g. Lyon and Ziegler 2009, Risse-Buhl et al. 2012, Kuehn et al. 2014, Hotchkiss and Hall 2015). The degradation of natural DOC by bacteria has been inferred from bacterial respiration (e.g. Cory et al. 2014, Mineau et al. 2016, Demars 2018), and many bacterial production estimates have been published (e.g. Fischer and Pusch 2001, Fukuda et al. 2006). The interaction strength between autotrophs and bacteria has been inferred statistically along nutrient gradients (e.g. Carr et al. 2005, Scott et al. 2008), but this is the first study, to our knowledge, quantifying the reciprocal carbon subsidies between autotrophs and bacteria in streams, notably bacterial CO_2_ use by autotrophs. This recycling of CO_2_ in turbulent headwater streams is possible within the intricate matrix of the biofilm (e.g. Kamjunke et al. 2015, Battin et al. 2016) or an algal mat, as when the CO_2_ is released in the water column it is very quickly degassed to the atmosphere (here 5-15 min, Demars 2018). It is important to note that the reciprocal carbon subsidies were identified under low flow conditions when stream water was hydrologically disconnected from soil water. When the soils are hydrologically connected to the stream water, and supply high fluxes of DOC, bacterial respiration and production was shown to be virtually independent of primary production (Demars 2018).

Our flow food web under low flows (end of experiment) suggested a strong microbial loop (to autotrophs) in the control stream, as expected at the low end of the nutrient gradient (Scott et al. 2008, Lyon and Ziegler 2009). The addition of labile DOC weakened this microbial loop, as indicated by the difference in reciprocal subsidies between algae and bacteria in the control and treatment (Fig. 10, 11), as predicted under nutrient limitation (Fig. 1). Bacterial production in the treatment reach relied more on sucrose than autotrophs and allochthonous organic matter, assuming a similar bacterial growth efficiency for all sources (range 5-20%), which is plausible under nutrient limitation (del Giorgio and Cole 1998) and corresponded to previous studies (e.g. Berggren et al. 2009, Fasching et al. 2014, Berggren and del Giorgio 2015).

The bacterial CO_2_ flux to autotrophs assumed its δ^13^C signature was the same as that of the bacteria, but the proportion of C sources used in bacterial production may be different to the proportion of C sources respired by bacteria. The controlled experiment allowed to calculate the bacterial respiration of the added sucrose as 2.16 g C m^−2^ day^−1^ over the three weeks of sucrose addition. This represented 82% (53-184%) of the estimated treatment bacterial respiration (2.64 ±1.45 g C m^−2^ day^−1^, Table S7) and the proportion of sucrose in bacteria was estimated at 51 ±7 % (Fig. S2). So the estimates were still within measurement errors.

The autotrophs did not include bryophytes in our flow food web calculations because their contribution to primary production was thought to be negligible over the few weeks of the experiment, and particularly towards the end of the experiment (on which data the flow food webs were based) when filamentous green algae were covering bryophytes. The lack of bryophyte growth under very low phosphorus concentrations (here 2-4 µg P L^−1^ of soluble reactive P) combined with shading by epiphytes has been well documented (e.g. Finlay and Bowden 1994). This may also explain the lack of changes in bryophyte C:N:P stoichiometry (Fig. 7).

### Boom and bust: role of nutrients

Gross primary productivity (GPP) appeared to be stimulated by the addition of sucrose but this was short lived, despite sustained light availability during the addition period (Fig. 4). Heterotrophic respiration of sucrose peaked after two weeks (three days after GPP) but crashed within days while the supply of sucrose was continuously flowing through the reach (with sucrose concentration increasing from 0.22 to 0.88 mg C L^−1^ with falling discharge). This was in sharp contrast to the peaks in respiration followed by a more sustained response of ecosystem (mostly heterotrophic) respiration to hydrological connectivity with soil water in the control reaches (Fig. 4, see Demars 2018). This boom and bust in the treated reach was likely due to nutrient limitation, mostly P according to the changes in filamentous green algae and periphyton C:N:P stoichiometry. Surprisingly, the shortfall of nutrients (N, P) was not compensated by faster nutrient cycling rates, as observed in previous studies with much higher labile DOC additions (e.g. Bernhardt and Likens 2002). This may be due to limitation of phosphate uptake by NH_4_ availability (long term median in both streams 7 µg N L^−1^) in streams relatively rich in nitrate (see Oviedo-Vargas et al. 2013). The higher primary productivity in the Cairn burn may partly result from higher P availability (4 µg P L^−1^) than in the control Birnie Burn (2 µg P L^−1^), despite lower nitrate availability (90 µg N L^−1^ *versus* 180 µg N L^−1^, respectively) – Table S3. These different nutrient supply rates may reflect the legacy of past experimental amendments (Ca, N, P, K) in 33 ha of the Cairn burn catchment aimed to increase grassland productivity in the late 1970s and early 1980s (Hill Farming Research Organisation 1983).

### Metabolic balance

The metabolic balance (or net ecosystem production) was responsible for a quarter of CO_2_ emissions in the control and increased from 6 to 12 % in the treatment reach during sucrose addition. CO_2_ emissions from these streams were therefore largely dominated by soil CO_2_ derived from the mineralisation of soil organic matter, rather than rock weathering of Dalradian acid schist drifts (Demars 2018). The proportion of in-stream biotic emissions were comparable to continental scale studies (<14% in African rivers, Borges et al. 2015; 28% in north American rivers, Hotchkiss et al. 2015). It is important to note that ecosystem photosynthesis and community respiration can be decoupled at individual sites in long term studies with temporal changes in C supply (e.g. Roberts et al. 2007, Beaulieu et al. 2013, Demars 2018). Under summer low flows respiration activity is more constrained by gross primary production explaining the strong interaction between photosynthesis and respiration (e.g. Demars et al. 2016).

### Fate of carbon

Sucrose is very labile and is well known to promote the growth of filamentous bacteria *Sphaerotilus natans* (“sewage fungus”), even at relatively low concentrations (0.25-1.00 mg L^−1^) in a stable flow forested stream (Warren et al. 1964). This was not observed in this study, likely because the moorland streams studied here were more open and colonised by *Microspora*, a common genus of filamentous green algae in Scottish streams (Kinross et al. 1993). *Microspora* was able to uptake sucrose by osmotrophy (Wright and Hobbie 1966) but this accounted to only 10% of C uptake by *Microspora*. Overall daily uptake of sucrose by autotrophs represented 1.2% of the average daily flux of sucrose. e proportion of added labile carbon (sucrose) in the consumers varied widely between species (Table S5), as observed in previous studies (e.g. Hall 1995, Hall and Meyer 1998, Collins et al. 2016), but not across functional feeding groups (19-27%), except for filter feeders (64%). This is questioning the usefulness of functional feeding groups, as defined here, to construct food webs. Filter feeders and notably blackflies (Simulidae) have been shown to directly assimilate DOC, extracellular polysaccharides and colloidal particles (Couch et al. 1996, Hershey et al. 1996, Ciborowski et al. 1997, Wotton 2009). This can explain the high proportion of added sucrose in blackflies (81%). Since the densities of blackflies were low (average 88-231 individuals m^−2^), C flux from direct uptake by consumers (filter feeders) were extremely small. The mass of sugar retained by all consumers was only 292 ±107 mg C m^−2^, or about 25 g C for the treated stream reach, representing 0.2% of the sucrose flux over the three-week addition.

The treatment reach did not show the large peaks in respiration (outside the period of sucrose addition) that the control reaches showed when the catchment was hydrologically connected, as indicated by soil moisture (Fig. 4, Demars 2018). This is likely because the treatment reach is a more constrained reach largely disconnected from the land. It was initially chosen to avoid lateral inflows which were very small (2.3%) and mostly from a spring fed flush (i.e. groundwater), rather than seepage from organic and riparian soils known to stimulate bacterial activity (e.g. Brunke and Gonser 1997, Pusch et al. 1998). Interestingly the average bacterial respiration was similar in the control and treatment reaches over the three weeks of sucrose addition (Fig. 10), suggesting part of the organic matter respired in the control was relatively labile and comparable to the 0.5 mg C L^−1^ of added sucrose. The dynamic of this respiration within the three weeks was very different however, decreasing with the progressive loss of hydrological connectivity with soils in the control, and peaking after two weeks in the treatment (Fig. 4).

By and far, the largest quantity of carbon processed by bacteria was lost as CO_2_ emission. Heterotrophic respiration over the treated reach respired (on average) 35% of the added sucrose. Bacterial production averaged 2-10% of the sucrose flux. Heterotrophic respiration of natural allochthonous DOC was about 3% in the control stream and 2.2 % prior to sucrose addition in the treatment. The fact that sucrose was processed 10 times faster than natural DOC was well reflected by the shortening of the organic uptake length (Sw_OC_) and increased mineralisation velocity (*v_f_*-OC). This was not surprising (see e.g. Marcarelli et al. 2011, Mineau et al. 2016), but the rate of mineralisation of natural DOC was relatively high and, scaled up to the first order catchment, represented 23±11% of the annual DOC flux in the control reach of the Cairn burn (Demars 2018). The changes in organic uptake lengths in the control and the differences between control and treatment prior to sucrose addition can be explained by the loss of hydrological connectivity with soil water, as indicated by the changes in soil moisture, and the difference in lateral inflows between the control (10.7%) and the treatment (2.3%). The shortening of the organic uptake length (Sw_OC_) reflected more hydrological changes as indicated by the different direction of change in organic carbon uptake (v_f_OC) between the control and the treatment. In the control v_f_OC declined by 32% against a 52% decline in DOC supply. In contrast, v_f_OC increased by 43% in the treatment with the addition of labile carbon, despite a 55% fall in DOC supply (similar to the control). Overall the uptake lengths of natural DOC (i.e. excluding those influenced by sucrose addition) were longer (2.5-4.3 km) than the length of the first order streams studied here (1 km), and so a large part of the carbon is released to downstream ecosystems as previously observed (e.g. Wiegner et al. 2005), especially during time of loss of hydrological connectivity with the soil of the catchment (Demars 2018). The simple comparison between flow food webs (Fig. 10) suggest that the addition of labile carbon did not prime the bacterial use of natural allochthonous DOC.

### Choice of carbon for DOC addition

We initially considered to add natural DOC to the stream after isolating DOC using reverse osmosis (Sun et al. 1995; RealSoft Pro2S, US) but the product was too salty with high pH and high nutrient concentrations (Stutter and Cains 2016). While reverse osmosis may be combined with electrodialysis to avoid co-concentration of salt (Koprivnjak et al. 2006), the quantities needed for our whole-ecosystem experiment were simply too large. We also considered several commercial humate sources but rejected their use because of pH, solubility and nutrient issues. Sucrose derived from sugarcane (C4 carbon fixation) has a very distinctive carbon stable isotope signature compared to the autotrophs in temperate ecosystems (C3 carbon fixation). It offered the possibility to trace its fate through the food web, and even to identify the bacterial carbon pathways in the control streams. It turned out that sucrose was a more judicious choice than first thought because labile DOC (polysaccharide, amino acids) is likely driving the respiration of the studied streams at the land-water interface (Demars 2018), as found in bioreactors (e.g. Drake et al. 2015).

## Conclusions

Part of the carbon derived from allochthonous organic matter can feed the autotrophs via the CO_2_ produced by stream bacterial respiration, intermingling the green and brown webs. The interaction between autotrophs and bacteria shifted from mutualism to competition with carbon addition under nutrient limitation (N, P) increasing biotic CO_2_ emissions. Without nutrient limitation, mutualism could be reinforced by a positive feedback loop, maintaining the same biotic CO_2_ emissions. Even a small increase in labile dissolved organic carbon supply due to climate and land use change could have large effects on stream food web and biogeochemistry with implications for the global C cycle under stoichiometric constraints.

## Acknowledgments

We thank Carol Taylor and Helen Watson for managing the long-term monitoring, Yvonne Cook and Susan McIntyre for running water chemical analyses, Claire Abel for the phospholipid fatty acid extraction, Maureen Procee for running the compound specific isotope ratio analysis, Gillian Martin for preparing and running the samples for stable isotope ratio analysis, Glensaugh farm manager Donald Barrie for hosting BOLD and JLK during the experiment and facilitating our work, and Baptiste Marteau for help with macroinvertebrate identification and comments on the manuscript. This study was funded by the Scottish Government Rural and Environmental Science and Analytical Services (RESAS), with additional funding support as part of the UK Environmental Change Network (ECN), and NERC Macronutrient Cycles Program. The writing up was partly funded by the Norwegian Institute for Water Research (NIVA). The authors acknowledge the provision of data forming part of the ECN wide dataset, https://catalogue.ceh.ac.uk/documents/456c24dd-0fe8-46c0-8ba5-855c001bc05f.

